# Generative adversarial networks for reconstructing natural images from brain activity

**DOI:** 10.1101/226688

**Authors:** K. Seeliger, U. Güçlü, L. Ambrogioni, Y. Güçlütürk, M. A. J. van Gerven

## Abstract

We explore a method for reconstructing visual stimuli from brain activity. Using large databases of natural images we trained a deep convolutional generative adversarial network capable of generating gray scale photos, similar to stimUli presented during two functional magnetic resonance imaging experiments. Using a linear model we learned to predict the generative model’s latent space from measured brain activity. The objective was to create an image similar to the presented stimulus image through the previously trained generator. Using this approach we were able to reconstruct structural and some semantic features of a proportion of the natural images sets. A behavioral test showed that subjects were capable of identifying a reconstruction of the original stimulus in 67.2% and 66.4% of the cases in a pairwise comparison for the two natural image datasets respectively. our approach does not require end-to-end training of a large generative model on limited neuroimaging data. Rapid advances in generative modeling promise further improvements in reconstruction performance.

## 1 Introduction

Since the advent of functional magnetic resonance imaging (fMRI), numerous new research directions that leverage its exceptional spatial resolution, leading to classifiable brain activity patterns, have been explored (Haynes, 2015). New approaches to decoding specific brain states have demonstrated the benefits of pattern-based fMRI analysis. Pattern-based decoding from the visual system has shown that it is possible to decode edge orientation (Kamitani and Tong, 2005), perceived categories of both static and dynamic stimuli (Haxby, 2001; huth et al., 2016), Up to identifying a specific stimulus image (Kay et al., 2008) and generically identifying new categories from image descriptors predicted from brain activity (Horikawa and Kamitani, 2017).

Here we focus on an advanced problem in brain decoding, which is reconstructing a perceived (natural) visual stimulus. The reconstruction problem is demanding since the set of possible stimuli is effectively infinite. A functioning reconstruction system may however prove highly Usefiil for neuroscience, for instance for studying synesthesia and optical illusions; or drive explorative insight into visual cortex activity when controlled experimental setups are difficult – such as during imagery or visual hallucinations. This problem has been explored at different spatial scales (e.g. invasively at the cellular level (Chang and Tsao, 2017)) and in different regions of the visual system (e.g. in the LGN (Stanley et al., 1999) and in the retina (Parthasarathy et al., 2017)). In this manuscript we discuss a new method for reconstruction from brain activity measured with fMRI. This approach was pioneered by Thirion et al. (2006), who reconstructed dot patterns with rotating Gabors from perception and imagery. Miyawaki et al. (2008) Used binary 10 × 10 images as stimuli and demonstrated the possibility of decoding pixels independently from each other, reconstructing arbitrary new images with this basis set. Naselaris et al. (2009) introduced a combination of encoding brain activity with structural and semantic features, as well as a Bayesian framework to identify the most likely stimulus image from a very large image database given the brain activity. Combining the most likely stimuli from a database leads to effective reconstructions, with (Nishimoto et al., 2011) being the most impressive example to date. These approaches were farther developed. Examples are enhancing decoding Using feature sets learned with independent component analysis (Güçlü and van Gerven, 2013) and accurate reconstruction of handwritten characters Using stimulus domain priors and a linear model for predicting brain activity (Schoenmakers et al., 2013, 2015). The most recent entries in the reconstruction domain make Use of promising new developments in generative image models. DU et al. (2017) Used Bayesian inference to derive missing latent variables, and effectively reconstruct handwritten digits and 10 × 10 binary images. Finally, the idea of reconstruction based on adversarial training has been Used for reconstructing face photos from fMRI with much low-level and abstract detail by learning to encode to and decode from a learned latent space for faces (Güçlütürk et al., 2017).

In this work we expand on the idea of Using adversarial training for reconstruction, but explore the capabilities of reconstructing arbitrary natural images Using generative adversarial networks (GANs, (Goodfellow et al., 2014)). We train a deep convolutional generative adversarial network (DCGAN, (Radford et al., 2015)) separately on large image data sets and let it learn the latent space in an Unsupervised manner. This DCGAN is Used to generate arbitrary images from the stimulus domain (handwritten characters or natural gray scale images). Keeping this DCGAN fixed, we learn to predict the latent space of the generator based on the fMRI BOLD signal in response to a presented stimulus. The objective of the predictive model is achieving high similarity between the generated and the original image in the image domain. The image domain losses that are Used to train the predictive model are derived with a complex loss function. We show that this approach is capable of generating reasonable reconstructions from fMRI data for the given stimulus domains. The method presented here is not limited to fMRI, but can be applied equally to other pattern-like responses measured for static images, which include calcium imaging or multielectrode arrays. Given suitable generative models for dynamic stimuli (such as video and audio), it may also be possible to transfer the method to other stimulus modalities.

## 2 Methods

### 2.1 Functional MRI data sets

We made Use of three publicly available fMRI data sets originally acquired for experiments related to identifying stimulus images and categories or reconstruction of perception. In the following we briefly list their properties. Extensive descriptions of recording details and methods can be found in the original publications.

#### 2.1.1 Handwritten characters

We Used this dataset (referred to as `BRAINS` dataset) to test our method in a simpler, restricted domain. Three subjects were presented with grayscale images of 360 examples of six handwritten characters (B, R, A, I, N and S). Images were taken from data published in (Van der Maaten, 2009; Schomaker and Vuurpijl, 2000). stimuli were presented foveally with fixation in a 3T fMRI experiment (TR=1.74 s, voxel size=2 mm^3^). The images were shown for 1 s at 9 × 9° of visual angle, flashed at approximately 3 Hz. The characters were repeated twice, and responses were averaged. The original studies reconstructed handwritten characters Using a linear decoding approach (Schoenmakers et al., 2013) and Gaussian mixture models (Schoenmakers et al., 2015). We made Use of the preprocessed data from V1 and V2 available in the BRAINS dataset and Used the original train / test set split (290 and 70 class-balanced characters respectively). The voxel response (*β*) per image was estimated in the original study, with a GLM and a canonical HRF, and we Used the preprocessed result of their method. The dataset can be downloaded from www.artcogsys.com.

#### 2.1.2 Masked natural images

Three subjects saw natural gray scale images with a circular mask, taken from different sources (the commercial Corel Stock Photo Libraries from Corel Corporation, and the Berkeley Segmentation Dataset) at 20 × 20° of the visual field with fixation. The dataset and experiments were described in (Kay et al., 2008) and (Naselaris et al., 2009). The training set consisted of 1750 images, presented twice and averaged. The test set consisted of 120 images, presented 13 times. Images were presented for 1 s and flashed at approximately 3 Hz. Data was acquired in a 4T scanner (TR=1 s, voxel size=2 × 2 × 2.5 mm^3^). The dataset is available on www.crcns.org Under the identifier `vim-1`^1^, which is also how we refer to it in this manuscript. We obtained a version of the dataset with Updated preprocessing for all three subjects from the author via personal communication. The peak voxel responses (as *β*) per image were estimated in the original study, with a voxel-wise HRF, and we Used the result of their method here. The advantage of this dataset for this study is the relatively high amount of data and the variety of high-quality photo stimuli.

#### 2.1.3 Natural object photos

This dataset was originally recorded for (Horikawa and Kamitani, 2017), and is referred to as `Generic Object Decoding` dataset. Five subjects were presented with square colour images from 150 categories from the ImageNet database (Deng et al., 2009). We converted the stimulus images to gray scale and applied a similar mild contrast enhancement as in (Kay et al., 2008) instead of Using the fUll color stimuli for reconstruction^2^. We also Used the original train / test set split. The training set consisted of 8 images from each category and was presented once, totaling 1200 presentations. The test set recording consisted of presenting single images of 50 categories (not contained in the training set) 35 times each, and averaging this activity. The image-wise response was estimated by averaging over the 9 s image presentation period. The data can be obtained from www.brainliner.jp^3^. Next to having recordings of five subjects one advantage of this dataset is the long stimulation time of 9 s (at 2 Hz flashing) per image, resulting in a high signal-to-noise ratio (SNR). All images were presented at 12 × 12° of visual angle, with fixation, in a 3T scanner (TR=3 s, voxel size= 3 mm^3^).

The data of the individual subjects of all datasets were mapped to a common representational space based on hyperalignment (Haxby et al., 2011) Using PyMVPA^4^ (Hanke et al., 2009). Hyper-aligned data was averaged across subjects such as to obtain data for a single hyperaligned subject with improved SNR^5^. Details about this procedure and the original voxel dimensions per subject, and resulting common representational space dimensions can be found in Section A.2. After hyperalignment, the dimensionality of the feature (voxel activity) space was reduced by applying principal component analysis (PCA, including demeaning) so that 99% (`BRAINS` leading to 248 dimensions, `Generic Object Decoding` leading to 1078 dimensions) or 90% (`vim-1`, dUe to its much larger original voxel dimension, leading to 2101 dimensions) were preserved. Hyper-alignment, PCA and statistical parameters (e.g. mean values) were computed on the training sets and applied on the training and the separate test set. For these additional preprocessing steps we Used the single trial data for `vim-1` and `Generic Object Decoding`, as the different averaging strategies changed SNR between train and test. For `BRAINS` we Used the provided data averages over two trials as there was no such difference between the train and test recordings.

### 2.2 Generative Adversarial Networks

Generative Adversarial Networks (GANs, (Goodfellow et al., 2014)) learn to synthesize elements of a target distribution *p_data_* (e.g. images of natural scenes) by letting two neural networks compete. Their results tend to have photo-realistic qualities. The Generator network (*G*) takes an n-dimensional random sample from a predefined distribution – conventionally called latent space **z** – and attempts to create an example *G***(z)** from the target distribution, with **z** as initial values. In the case of images and deep convolutional GANs (DCGANs), introduced in (Radford et al., 2015), this is realized across a series of deconvolutional layers. The Discriminator network (*D*) takes a generated or real example as input and has to make the binary decision whether the input is real or generated, which would result in the output 1 or 0, respectively. In the discriminator, DCGANs use a series of convolutional layers with a binary output. See Figure 1 for an illustration. This competition process is expressed as a zero-sum game in the following loss term:

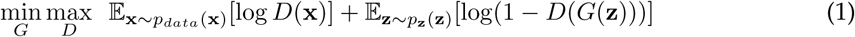

where **x** is a real image and *G***(z)** a generated image. During training, various – but certainly not all – GAN variants learn to impose structure on the latent space. Learning this structure and the learning procedure itself is a form of unsupervised learning. The algorithm we use was introduced and popularized for image synthesis by (Goodfellow et al., 2014). Creswell et al. (2017) is a recommended comprehensive review and discussion of various recent GAN approaches.

**Figure 1:**
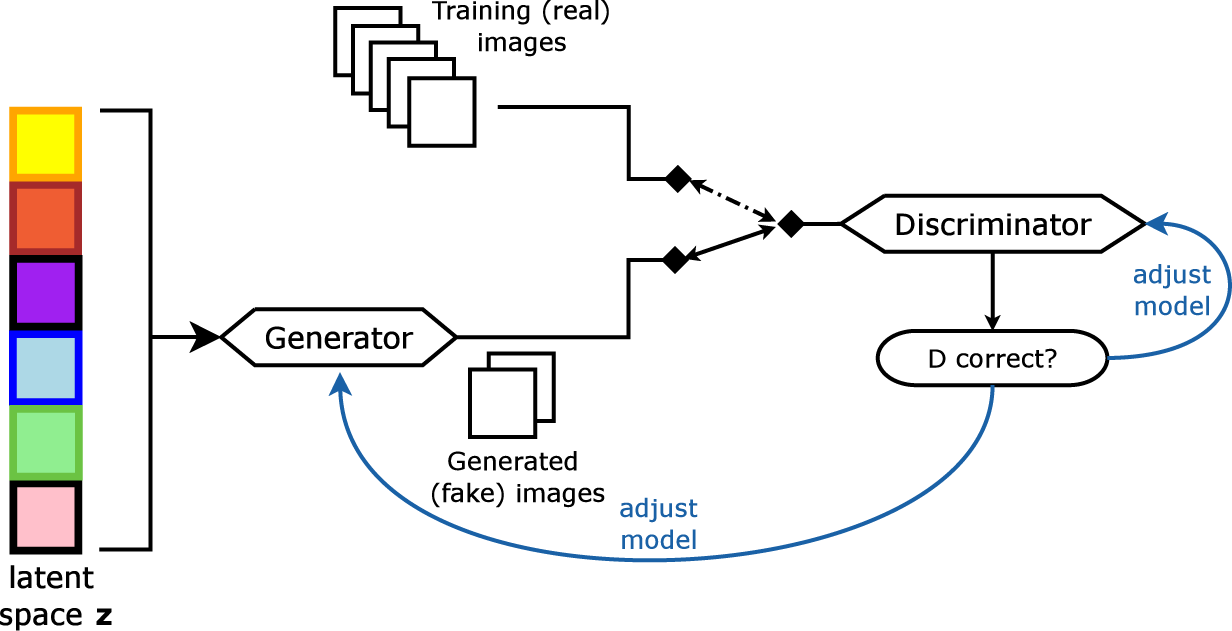
Generative adversarial networks. A generator network (G) learns to model a given distribution *p_data_* via feedback from a discriminator network (D). D learns to discriminate between images coming from the real distribution and images from the generator.

For this work we used a DCGAN architecture that implements architectural improvements suggested in (Radford et al., 2015) and (Salimans et al., 2016). We based the model on a publicly available framework and implementation (`musyoku`, 2017). This generative model is merely a module of the method and can be replaced by any better-performing advanced deterministic generator. Details of the architecture and the training parameters are described in section A.1.

We trained the same DCGAN architecture separately for each dataset, for approximately 300 iterations through all training images. Figure 2 shows examples from the `vim-1` training set, and randomly generated examples from a DCGAN trained on this data. The network seems to have learned the contrast properties of the `vim-1` stimulus set, and seems to have acquired the ability to create complex image content. As we selected these random example images manually they reflect our preference for semantically meaningful content. Yet, as with most GAN architectures, much of what is created is rather abstract and can not be interpreted. The handwritten character GAN in contrast learned to create primarily meaningful new examples of the reduced handwritten character set. We noticed that it rarely generated B examples though. So the DCGAN architecture we are using potentially suffers from a form of the so far unsolved problem of mode collapse.

**Figure 2:**
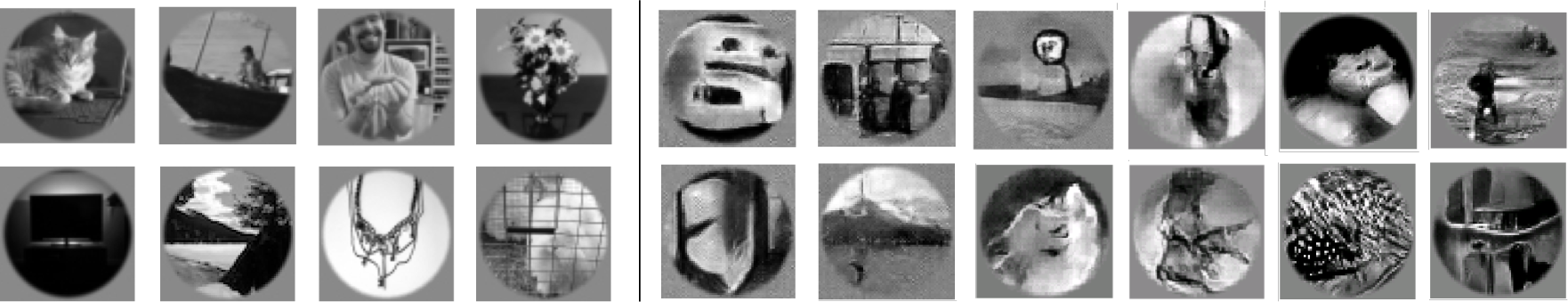
Training a DCGAN for naturalistic vim1-like gray scale images. Left: Examples from the gray scale natural image domain DCGAN training set (gray scale MS COCO or ImageNet; masked). Right: Examples of images randomly generated by a trained natural image DCGAN.

We checked whether the expressive power of the chosen DCGAN is sufficient for reconstructing natural stimuli from the experiments by overfitting the model predicting **z** from BOLD data on the training data. For this we used a multi-layer perceptron (MLP) instead of the linear regression approach outlined in the following section 2.3. In Figure 3 we show training set reconstructions on `vim-1` from such an overfitted model. These examples can also be seen as an upper limit of the accuracy that can be expected with the DCGAN architecture used here. It is obvious that especially broad high-contrast boundaries can be reconstructed, but the natural images DCGAN also seems to capture patterns, luminance, luminance gradients and some of the semantic content (e.g. landscapes) that are in the stimulus set. We thus can state that the natural image DCGAN reflects the reconstruction target sufficiently. We assume but can not verify that semantic content can be reproduced if structural properties of the image restrict the semantic space. For instance, landscape photos frequently feature a horizontal bar across the whole image.

**Figure 3:**
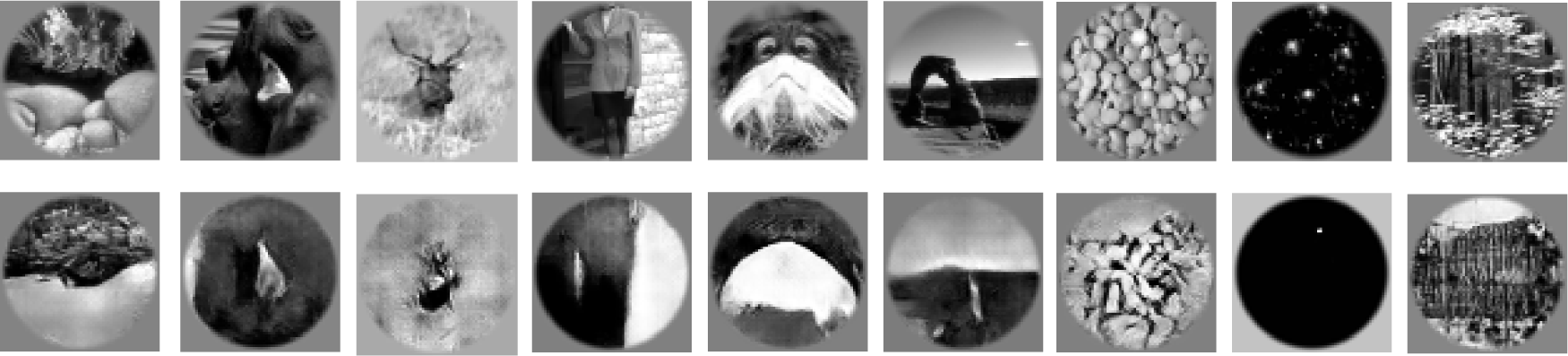
The natural images GAN captures the vim-1 training stimuli. We overfitted the model on random training set images to demonstrate that the latent space the GAN has learned is powerful enough to capture and regenerate the variety of the `vim-1` stimulus images satisfactorily. The top row shows the original stimulus image, the bottom row the overfitted reconstruction.

### 2.3 Latent space estimation from BOLD data

We fixed the trained DCGANs and attempted to predict the latent space **z** ∈ [−1, 1]^50^ that reproduces the correct image directly, with the estimated BOLD data per image as the independent variable. The loss for this model was gathered in image space with a complex multi-component loss function that compares the real and the reconstructed images with pixel data and perceptual features learned by a convolutional neural network. The linear regression model was implemented as an approximating neural network with one weight layer. The procedure is illustrated in Figure 4. For weight optimization we used the Adam optimizer (Kingma and Ba, 2014), again with default parameters. We applied a weak L2-regularization on the weights (λ_*L*2_ = 0.01). The same normalization that we used as a boundary on **z** when training the GAN was applied to the predicted **z** after applying the linear regression weights. The model was trained for 300 epochs on BRAINS and `vim-1`, and for only 100 epochs on Horikawa as the model seemed to learn this data set faster (and could overfit if trained much longer).

**Figure 4:**
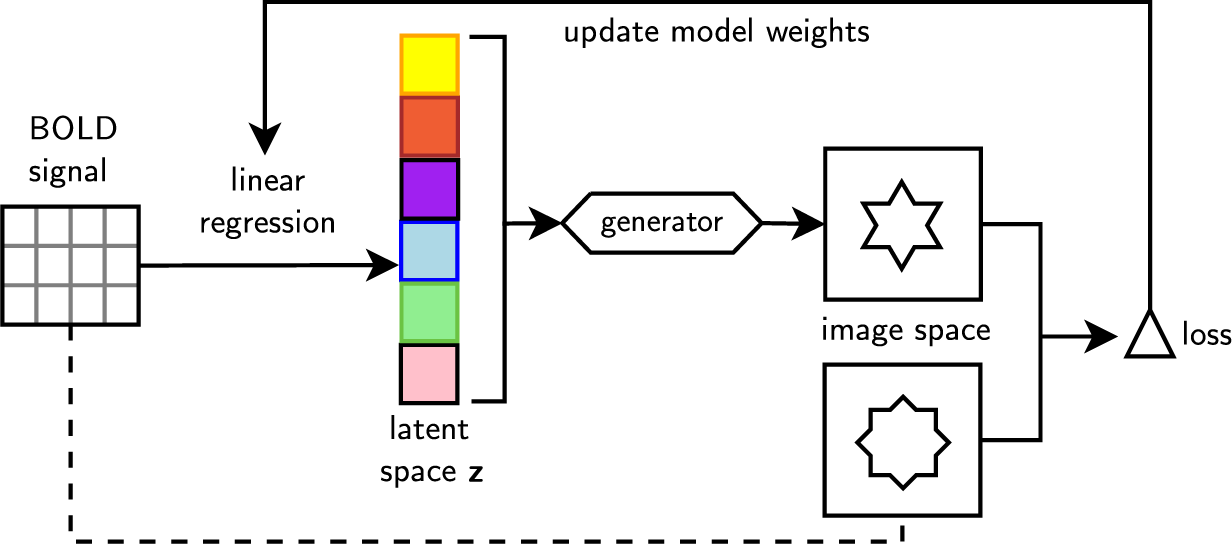
Predicting z from BOLD data with a complex loss function in image space between the reconstructed image and the image actually shown in the experiment. We make use of the DCGAN generator, which is pretrained for the necessary stimulus domain and not updated further during reconstruction model training.

To compute the loss for training, we passed the predicted latent vector **z** for every batch (using a mini batch size of 3) through generator network *G* of the previously trained DCGAN. The image produced by the generator *G***(z)** (reconstructed) was compared to the image **x** actually shown in the experiment with a complex image loss function. This loss function is a weighted sum of the following components (formulated as an average over mini batches):

#### Mean absolute error on pixels

The 64 × 64 images were downsized by 10% with bilinear interpolation. This is merely for avoiding the blurring effect of the MSE observed in autoencoders. Then mean absolute error (MAE^6^) was calculated between them to obtain the pixel loss *l_px_*:

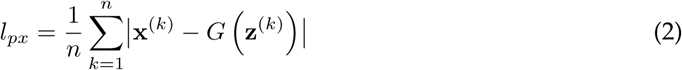

where *k* ranges over images in the image batch of length *n* The pixel loss optimizes towards matching local luminance values between reconstruction and the stimulus images.

#### Feature losses

Weights learned by convolutional neural networks on natural image data present a set of low- and intermediate level feature detectors useful for various applications. In methods such as autoencoders a feature-based loss seems to lead to higher perceptual similarity, whereas using pixel-wise loss functions such as MSE leads to blurry results (Johnson et al., 2016). We trained a variant of the AlexNet neural network to obtain suitable feature detectors (Krizhevsky et al., 2012).

Here, the perceptual *features* of an image are defined as the outcome of passing an image through a trained multi-layer neural network and obtaining its hidden unit activity matrix at certain layers in its hierarchy. As an example, many convolutional neural networks trained for object recognition in natural images learn a series of oriented Gabor filters in their first layer. After passing an image through the network and obtaining its activity in this first layer, we would have a matrix indicating local similarities to these Gabor filters (similar to the idea of receptive fields in neuroscience). This matrix is called *feature matrix*. Feature activations were gathered from the feature matching network before any nonlinearities or local response optimization.

For computing the loss between *G***(z)** and **x** on the basis of features from the trained AlexNet hierarchy (the *feature matching network*) we used the following loss component: The feature activation matrices of the real stimulus images for each layer *L*, denoted *ϕ*_*L*_(x) were transformed to binary representations *ϕ*_*L*,*b*_(**x**) by applying a threshold of 1.0. The second component of the reconstruction loss function is then *feature magnitude loss l_f,m_*, which equates mean squared error computed between *ϕ*_*L*_**(x)** and *ϕ*_*L*_(*G***(z))** on feature map elements that met the binarization threshold in the original stimulus image (1 in ϕ_*L*,*b*_(x)):

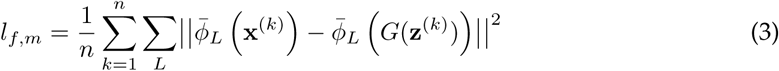

where 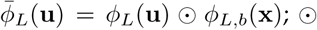 denoting element-wise multiplication. This is merely a formalization for using *ϕ_L,b_* (i.e. the binary matrix of feature matrix entries whose activation passed the threshold) as a binary mask. This loss term ensures that the activation magnitudes (saliency) of the various perceptual features stay similar to the real image and avoids some of the fallacies of using MSE directly between feature matrices^7^.

We used layers `convl, conv3` and `conv4` for feature matching as these represent universal low-level features and simple patterns in the AlexNet architecture. The highest layers of AlexNet may otherwise represent sparse semantic properties. Furthermore matching the final layers may also drive the reconstruction towards the limited set of categories learned by AlexNet. From `convl` we also collected *l*_*f*,*m*_ on negative feature activations, using the threshold 1.0 on their absolute representation, as they are collected via meaningful convolutions in pixel space. Negative feature activations for higher layers are likely meaningless as they are not used during training.

The complete loss function is then given by adding the terms with a weight:

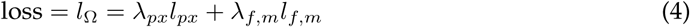

where we chose *λ*_*px*_ = 100.0 and *λ*_*f,m*_ = 1.0 for all three datasets. These weights, the set of layers used for feature matching and the feature threshold were determined via optimizing on the training set of `BRAINS`. Optimization specifically for each data set may improve the results further, however a cross-validated hyperparameter search would require more data than we had available. Also, these loss magnitudes are likely specific for the chosen DCGAN, the stimulus data set and the feature matching network and need to be redetermined when applying this method for a different experiment.

The gradients of the linear model weights w are estimated as follows:

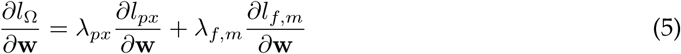

Remember that the reconstructed image *G(z)* is associated to w in the following way:

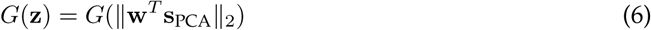

**s**_PCA_ is the BOLD signal represented by its principal components. Thus the gradients are estimated by backpropagating through the specific loss functions (mean absolute error for *l_px_*, or through the feature extraction network *ϕ*_*L*_ for *l*_*f*,*m*_), the generator *G* and the norm. Implementation-wise this backpropagation procedure relies on automatic differentiation techniques as implemented in all common neural networks frameworks.

#### Feature matching networks

Feature matching requires a universal set of image descriptors that frequently occur in our chosen natural images *p_data_*. To obtain these descriptors we trained a variant of AlexNet (Krizhevsky et al., 2012), with one input channel and 5 × 5 kernels in the first (conv1) layer on the 64 × 64 grayscale ImageNet data described before. The model was trained towards classifying the standard set of ImageNet categories. We used this network for `vim-1` and `Generic Object Decoding`, ignoring potential redundancy of features extracted from the mask in the former. For the `BRAINS` data set we again trained an AlexNet architecture. In this case we trained on all 40,000 examples of 36 handwritten digit and character classes from (Van der Maaten, 2009) and (Schomaker and Vuurpijl, 2000) in order to obtain a universal set of image descriptors for the handwriting domain.

#### Reconstruction variability

One inherent disadvantage of training models with random components, such as randomly initialized weights or stochastic gradient descent (e.g. neural networks) is the variability of the results, due to different local minima the model will converge to. Furthermore, in the case of GANs small shifts in the predicted latent space can result in well-perceivable changes in the generated image. We observed this behaviour, which resulted in the model finding different ways to reconstruct certain images, reconstructing different features of images, or not finding a recognizable reconstruction at all for an image that could be reconstructed in previous models. This variability is demonstrated in Figure 8. We attempted to counteract these effects when obtaining final reconstructions with a simple ensemble model: We averaged the predicted **z** over 15 independent training runs, normalizing **z** to the unit hypersphere (see Section A.1 for normalization details) again after this.

The feature matching networks, the natural images GAN and the predictive model for **z** have all been implemented in the `Chainer` framework for neural networks (Tokui et al., 2015)^8^.

## 3 Results

Using the outlined methods and parameters we obtained a set of validation set reconstructions for each data set, out of which we show examples of reconstructed images and failure cases in the following. We proceeded with a quantitative behavioural evaluation of overall recognizability on these sets.

### 3.1 Reconstruction examples

#### 3.1.1 Sample reconstructions on BRAINS

Figure 5 demonstrates that the method can lead to accurate reconstructions when the DCGAN is restricted to few handwritten characters classes. Despite a small training set of 290 BOLD images of V1 and V2, the correct handwritten character is reconstructed in 54% of the cases (determined via human rating; chance level: 17%). Successful reconstructions demonstrate that the model is also capable of reconstructing structural features such as position and curvature of lines. When character classes could not be reconstructed frequently such structural similaritiesremained. As mentioned before, the underlying handwritten character DCGAN had difficulties generating examples of B, and the reconstruction model also failed to reconstruct a B stimulus in 9 out of 12 cases.

**Figure 5:**
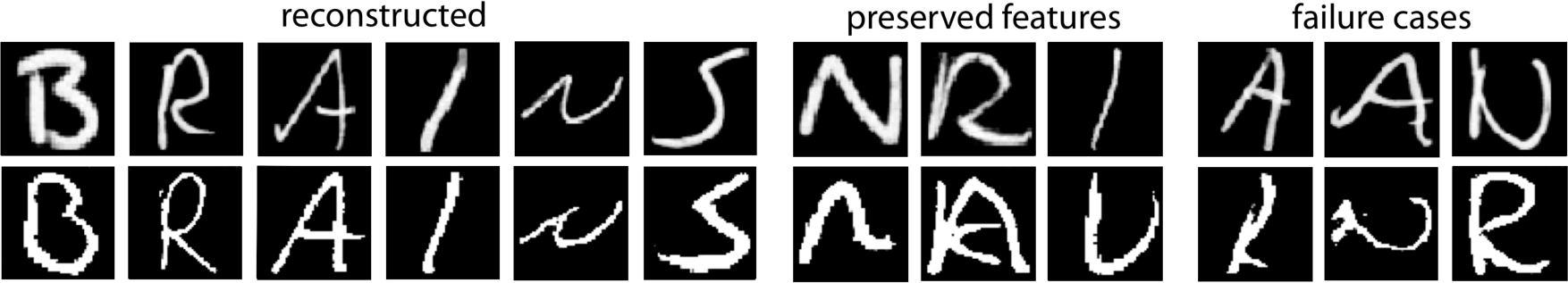
Reconstruction examples for handwritten characters. Top row: Presented stimuli. Bottom row: Reconstructions from BOLD activity.

#### 3.1.2 Sample reconstructions on vim-1

Figure 6 contains reconstruction examples for natural grayscale stimuli from the `vim-1` dataset. At 1820 training examples it was the largest training data set we used. In reconstructions that were sufficiently accurate to be identifiable in the behavioural tests, contrast differences appear to be the most likely image feature preserved. Salient pattern information also remained intact. In some reconstructions luminance information is lost, while structural features remain. A horizontal contour line across the image appears to lead the model into generating a landscape image, however not in every case.

**Figure 6:**
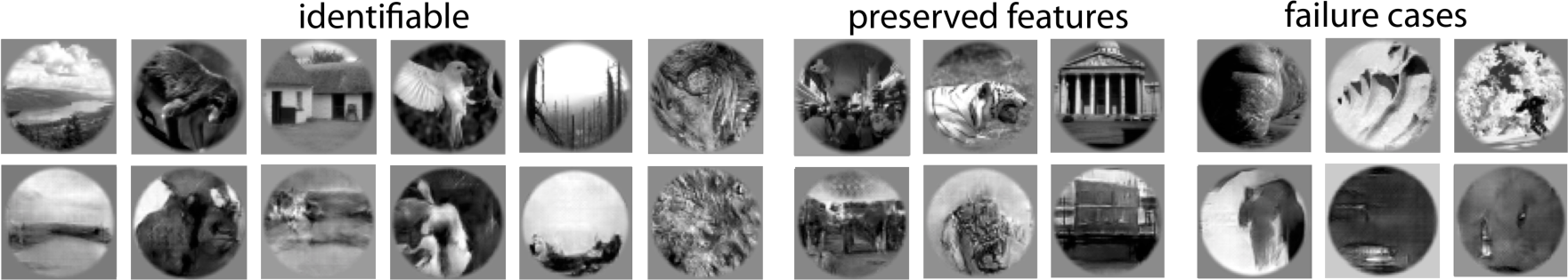
Reconstruction of natural grayscale images (vim-1). Top row: Presented images. Bottom row: Reconstructions from BOLD activity. Images in the *identifiable* category are reconstruction examples correctly assigned in no less than 8 of the 10 behavioural comparisons.

#### 3.1.3 Sample reconstructions on Generic Object Decoding

The `Generic Object Decoding` dataset has just 1200 training examples, but high SNR due to long stimulation time. The stimuli are not masked, so overall more content needs to be reconstructed per image. Reconstruction examples can be seen in Figure 7. Overall the reconstructions again preserve salient contrasts, but turned out more blurry than the `vim-1` reconstructions. There are also more failure cases^9^.

**Figure 7:**
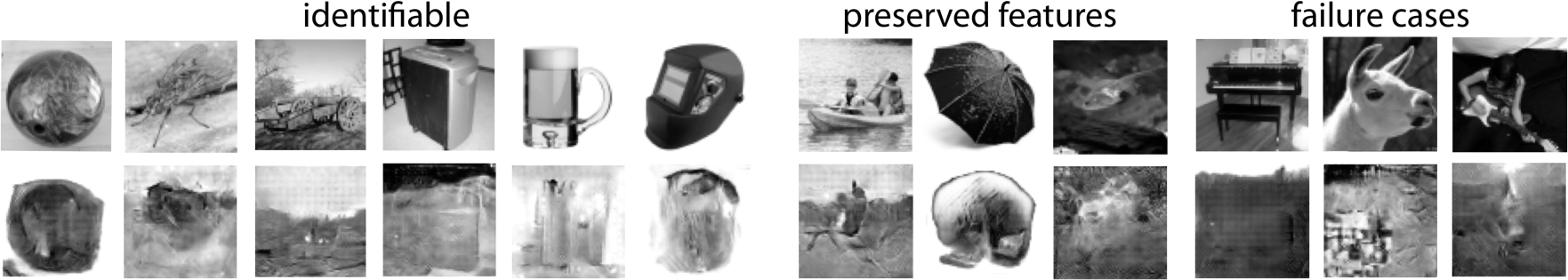
Reconstruction of natural grayscale images (Generic Object Decoding). Top row: Presented images. Bottom row: Reconstructions from BOLD activity. Images in the *identifiable* category are reconstruction examples correctly assigned in no less than 8 of the 10 behavioural comparisons.

#### 3.1.4 Reconstruction variability

When using different model parameters and loss weights, and to a lesser extent across different runs with the same parameters the model was often reconstructing the same images in different ways (for identifiable reconstructions). We described the potential cause of this behaviour further in Section 2.3 and demonstrate these effects in Figure 8. With the same parameter choices, using the ensemble model counteracts leftover variability.

**Figure 8:**
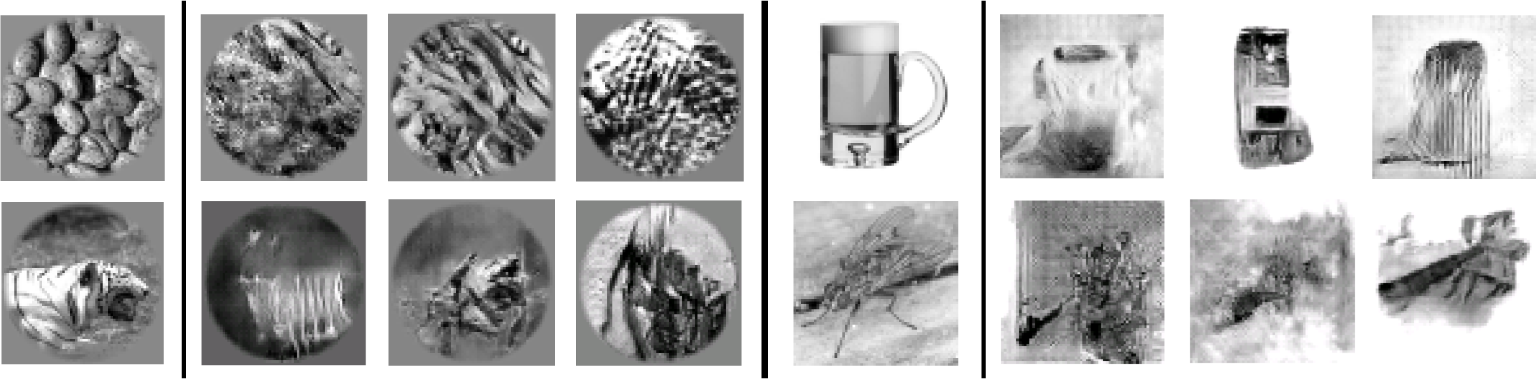
Reconstruction variability. Reconstructions vary when using different loss weights (λ) and layers, and to a lesser extent when running the model with the same settings multiple times. This is caused by the sensitivity of the **z** space and by different minima the model may find in the DCGAN. The image shows variants of reconstructions from different parameter settings and hyperparameter (layers, thresholds) choices when estimating the linear model with the same DCGAN.

### 3.2 Behavioural evaluation

A number of successful reconstructions of natural images have reversed luminance information or only slight or transformed structural similarity. Due to this, potential similarity measures such as the structural similarity index can not be applied as the comparison task is too complex in the natural images case. In order to obtain a quantitative measure of reconstruction similarity on each data set we instead made use of human perceptual systems.

We conducted a behavioural perceptual study on Amazon Mechanical Turk^10^. The advantages of this platform over common university subject pools for collecting human labeling and uncomplex behavioural scientific data have been discussed and demonstrated (Mason and Suri, 2012). Workers were presented with one original image from the validation sets and had to chose between the real and one randomly chosen different reconstruction taken from the same validation set. Each of these choices was one Human Intelligence Task (HIT) compensated with $ 0.01. As a preventive measure against fake completions and bots, workers had to hold the Masters status and have an approval rate of 95% or higher on the platform to qualify for our tasks. We repeated the procedure ten times for each of the validation set images in each data set, paired with a different randomly chosen reconstruction from the set in every HIT. To prevent gaming the task by memorizing the common reconstruction between two comparisons for the same image, it was made sure that no worker individual saw any image-reconstruction pair twice. Note that due to the nature of the system, it was also unlikely that any worker saw the full set of images of any category, though this did occur in few batches. Every HIT was presented once, i.e. we did not use the platform’s internal repetition mechanism for verification. Across all three validation sets we presented 2420 of these comparisons in total (due to different validation set sizes 500 for `Generic Object Decoding`, 1200 for `vim-1` and 720 for BRAINS). These 2420 comparisons were processed by 105 worker individuals in total.

Figure 9 shows worker performance for the validation images of the three data sets. The number of *correct decisions* denotes the total number of correct decisions across all HITs (comparisons). As there were ten such HITs per reconstruction it is slightly skewed both by failure and well-identifiable reconstructions, but is a better representation of overall undecidedness. The number of correct decisions *by image* on the other hand applies a majority vote over the ten decisions per image, representing the number of validation set images that could be correctly identified.

**Figure 9:**
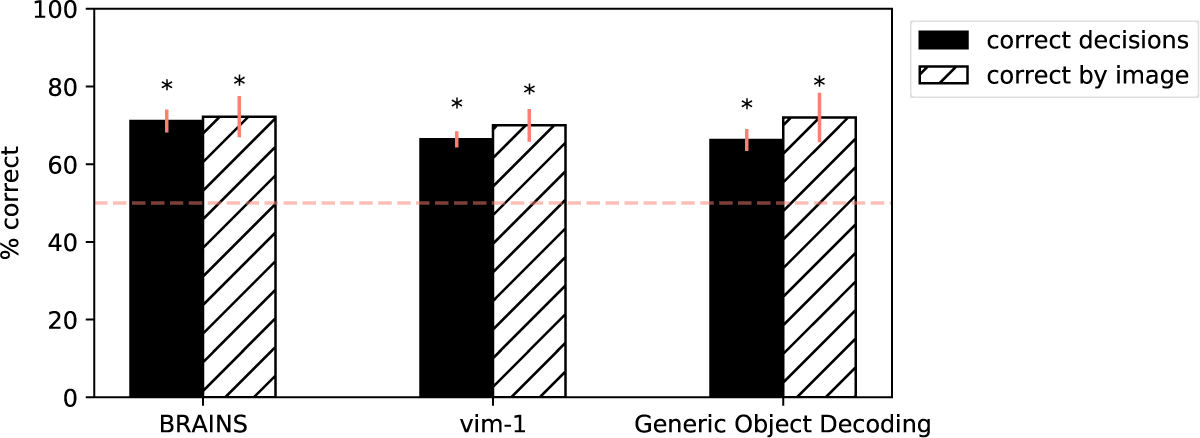
Correctly identified reconstructions in pairwise behavioural test. Mechanical Turk workers were presented with an original image and had to chose its reconstruction out of two. The error bars show the standard deviation, estimated via non-parametric bootstrapping.

All results were significantly different from random choices with *p* ≪ 0.01 based on a binomial test (see (Salzberg, 1997)). Although the BRAINS dataset model reconstructed the correct character class in a mere 54% of the validation set images, structural resemblance between the original characters and their reconstructions were still strong enough for 71.1% and 72.2% correct identifications based on overall and per-image decisions respectively. From the set of `vim-1` images workers correctly assigned 70% of the reconstructions, applying the majority vote per image. As for the overall number of correct decisions only 66.4% were correct, indicating that many reconstructions were still too crude to not be confused with a different random one. The `Generic Object Decoding` natural images stimulus set performed similar at 66.2% correct overall, and 72% if grouped by image. The errors shown in Figure 9 were estimated with non-parametric bootstrapping on the behavioural results, drawing 100,000 samples. Overall right decisions had standard deviations of 2.9%, 2% and 2.8% for the data sets `BRAINS`, `vim-1`, `Generic Object Decoding`. The majority vote per image-metric was estimated to have a standard deviation of 5.3%, 4.2% and 6.3% respectively.

Figure 10 shows how often individual images and their reconstructions were matched in the 10 pairwise comparisons per image. If at least 8 correct comparisons out of 10 means that an image was *identifiable* (see images of reconstructions), then 55.6% of the images in `BRAINS`, 43.3% in `vim-1` and 38% in `Generic Object Decoding` fell into this category.

**Figure 10:**
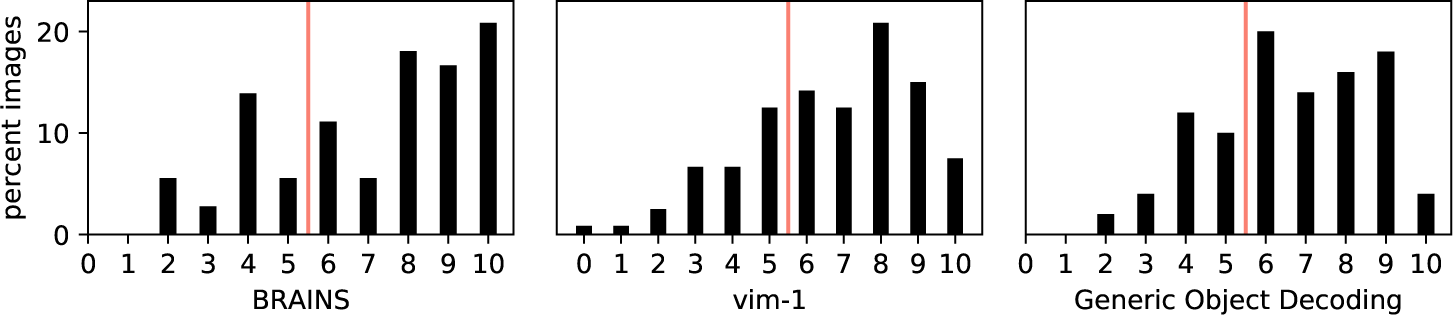
Histogram of number of correct decisions across validation set images. Shows how often a reconstruction was correctly assigned to the original image, out of 10 comparisons. The red line indicates the majority vote threshold in the *correct by image* metric.

### 3.3 Interpretability of the latent space in relation to brain representations

The fact that recognizable features of stimulus images could be reconstructed with a simple linear model indicates that the latent space represents properties that are also represented in brain activity. However the exact structure of the learned latent space is unknown, and most neural networks literature shows a selection of subjectively meaningful directions, but no overall analysis of properties.

In the following we follow a different, novel route for investigating what the latent space has learned. In research about natural image statistics the *distributions of image gradient magnitudes* in different images have been empirically (Simoncelli (1999)) and theoretically (Geusebroek and Smeulders (2002)) shown to follow Weibull distributions. These distributions are characteristic for different types of natural images. There are two parameters determining this distribution, shape (*γ*) and scale (*β*) (see Equation 7). For natural images *γ* and *β* are strongly correlated. Intuitively an increased *β* parameter means that a histogram of gradient strengths of an image would be shifted towards the right, which means that an image has more high contrast edges (see examples in Figure 11).

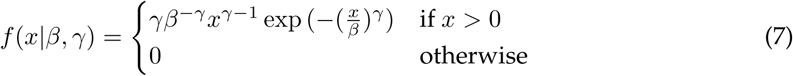

**Figure 11:**
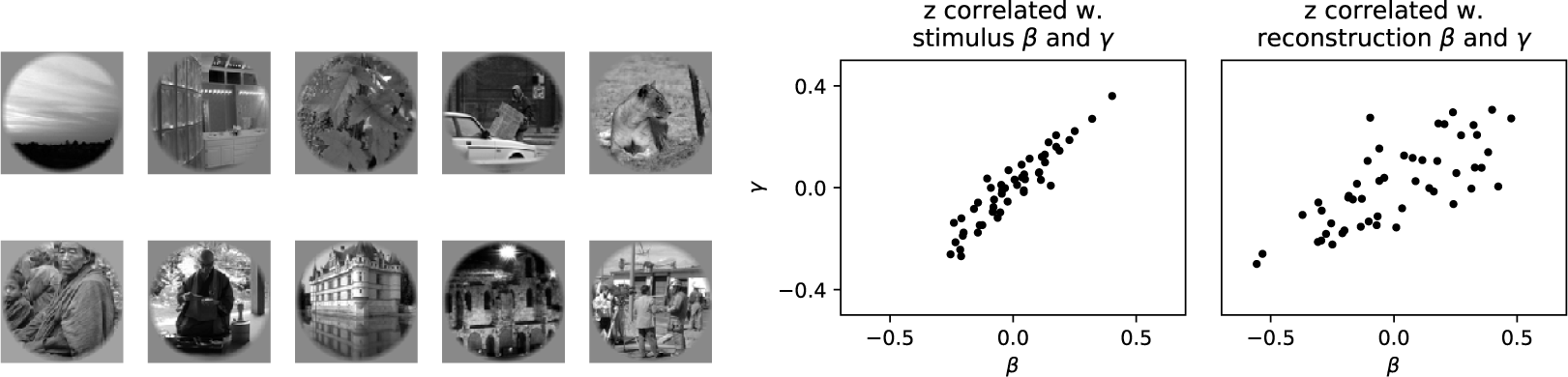
Latent space and Weibull image statistics. Left: Example images from `vim-1` for increasing *β* (row-wise from top left image to bottom right image). Right: Pearson correlation between each of the 50 dimensions of **z** and the shape and scale parameters that describe the distribution of edge magnitudes. Given for original images and for reconstructed images.

The shape and scale parameters have been shown to be strongly correlated with brain activity as measured in electroencephalography (EEG) (Scholte et al. (2009), demonstrating 71% explained variance of the early grand-average ERP signal). We used 3×3 vertical and horizontal Sobel operators to determine the gradient magnitudes *x* for each original and reconstructed image in the data set `vim-1`. Then we modeled the probability density function of the image gradient distribution with the 2-parameter Weibull distribution, determining the shape and scale parameters via maximum-likelihood estimation. We then proceeded to compute the Pearson correlation between the shape and scale parameters of an image (original or reconstructed) against each estimated latent dimension.

It turned out that several of the latent space dimensions estimated by our linear model show moderately high positive and negative correlations (up to 0.6) with these two natural image statistics descriptors (see Figure 11, right panel). The earlier findings show that these parameters can be linearly associated with EEG brain activity, and in Figure 12 we also see this property for our fMRI data, were primarily voxels in VI, V2 and V4 are correlated with the image statistics parameters in the `vim-1` dataset.

**Figure 12:**
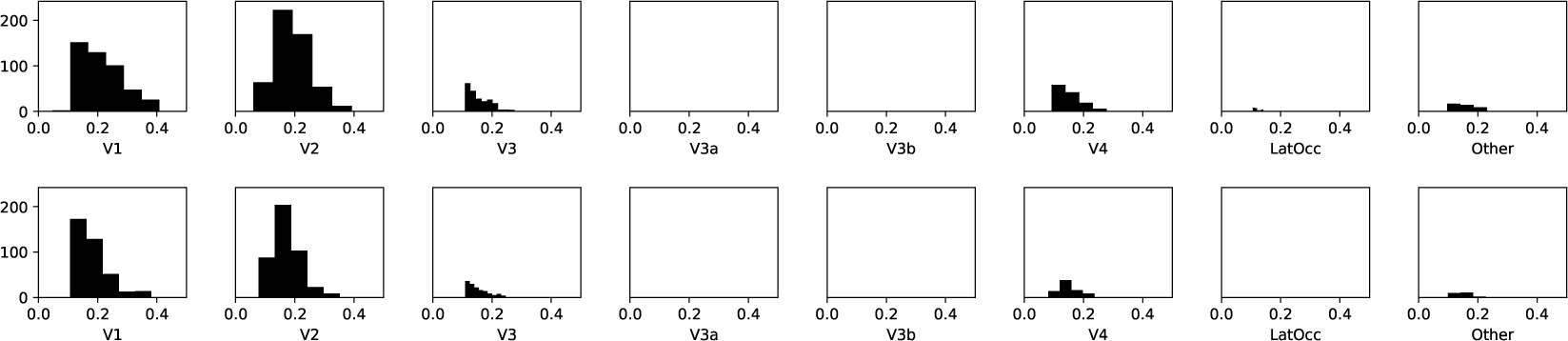
Correlations between vim-1 S1 data and image statistics. Histogram of significant voxel-wise correlations (*p* < 0.01) between voxel activity and Weibull statistics on S1 from the `vim-1` training set. *β* is shown at the top and *γ* at the bottom.

## 4 Discussion

We presented a new approach for reconstructing static visual stimuli with natural image characteristics from brain activity. We conducted a behavioural study indicating that the reconstructions this method achieves are sufficient for linking an original image to its reconstruction, even in the virtually infinite domain of natural images. Using a DCGAN generator as a pretrained natural image prior assures that the reconstruction results employ natural image statistics, preventing noisy images. An advantage of our method is that it does not require end-to-end training of a high number of complex neural network layers on usually limited neuroimaging data, using a pretrained DCGAN as a black box instead. Also, as the generative model is trained separately on a large separate data set there is less danger of overfitting it on the variance in the stimulus distribution.

The current research in the area of reconstruction, including this study, should be seen as purely explorative as the experimental requirements are high: Subjects must be highly attentive and cooperative as they need to passively view thousands of images (of which a part needs to be frequently repeated to get a cleaner signal for the actual reconstruction) and multiple sessions are needed to acquire sufficient training data. Alignment across sessions is merely a partly-solved problem. The state of research is far away from single-shot reconstruction systems.

In the constrained domain of handwritten characters the correct character class could not be reconstructed in all cases, but the accuracy was still well above chance level. The method could furthermore reconstruct sufficient structural detail so that the right reconstruction could often still be identified in the behavioural test, even when the reconstructed character was incorrect. In this simpler, restricted domain the model showed good performance with a very limited amount of training data. However the DCGAN had difficulties generating one of the six characters. Nevertheless results indicate that the method can be applied for reconstructing stimuli from such a limited domain, if made sure that all potential stimulus manifestations occur in the reconstruction model. In the handwritten characters dataset, in several cases not just the class but virtually all structural features were preserved.

As for any current reconstruction method, reliability of using them as a measuring tool is unclear. What we perceive as noise in the reconstruction may be a true prediction by a model reacting to sudden changes in visual activity. Similarly, clearly visible image properties may just be noise in the model or the data recording. One danger of the restriction to photorealistic reconstructions in our work is that an observer may be more likely to trust the reconstructions. Determining the reliability of a reconstruction with attention or SNR measurements may be one avenue for research that the subfield would benefit from as a whole.

In Figure 3 we demonstrated on training set examples that the DCGAN captured much of the variety of the `vim-1` natural grayscale image set. While we can state that our results present a step forward over previous models, the reconstruction quality and generalization performance on the validation set certainly leave much to be desired. It is possible that generalization performance could increase by merely adding much more training data. Although both our natural image data sets contained less than 2000 training images (an insufficient amount for many machine learning methods), due to the difficulties of neuroimaging experiments `vim-1` and `Generic Object Decoding` are already considered large experimental data sets within the community. Yet for reconstruction studies such as ours, massive amounts of high-SNR visual system data from single subjects may be necessary. Another related antagonist of the generalization capabilities of our approach is the noisy nature of neuroimaging data which does not agree with the sensitivity of the latent space. Subtle changes in **z** induced by noise in the brain data are capable of strongly changing the features of the reconstructed image. It is unclear whether this problem would remain in an experiment with much larger amounts of data with high quality.

Also, we achieved our results with a linear regression model. This linear relation promises interpretability of the relations between the latent space and brain representations, when the reconstruction model is powerful enough and sufficient care is applied. The fact that we could achieve our results with a basic linear model also means that any more advanced regression model iteratively trained with a complex image loss function could further improve over our results. The latent space **z** of DCGANs has been shown to be capable of learning structural features of the target domain faces in the original publication (Radford et al., 2015). While this motivated our model choice, it is unclear to which extent this applies to our natural images GANs. Also, until there is further investigation it should not be assumed that standard DCGANs learn a well-structured latent space.

Our loss function involves pixel luminance as well as edge and basic pattern information. We did not penalize the model on higher-order semantic information, e.g. by using the actual classi-ications or higher convolutional and fully-connected layers from our feature matching network. The set of training classes of a convolutional neural network is always restricted to a specific subset. Our reconstructions would thus remain restricted to a predefined set of classes and could not be called arbitrary. Yet finding a valid method of adding a semantic penalty to the results could be another way of strongly improving over our results.

Our natural image GAN is set up to approximate the distribution of all natural grayscale images. This is still a constraint on the set of images that can be reconstructed. It will not be possible to reconstruct non-natural image types, such as handwritten characters or comic scenes; unless the generative model can been trained to generate images with non-natural statistics as well. A GAN trained on a specific image database such as ImageNet or MS COCO will reflect their potentially unbalanced selection of categories (e.g. dog breeds), which presents another bias. In the current development state GANs can also fall into local minima where generated images show low variety. The generator can often fool the discriminator by learning a limited set of image types (modes of the image distribution) perfectly. This problem is known as mode *collapse*, and considered one of the more important issues to solve by the deep generative modeling community. One frequently explored remedy is providing binary categorical information along with **z** in a semi-supervised fashion. However, as mentioned before, such a discrete set of categories would present a severe limitation contradictory to our aim of reconstructing arbitrary images. A recent theoretical investigation into the training dynamics leading to mode collapse (along with a proposed solution) can be found in (Kodali et al., 2017). The recent (Bang and Shim, 2018) proposes a solution that does not suffer from image quality loss. There are other recent ideas addressing the problem with unsupervised methods (Unterthiner et al., 2017). If mode collapse remains unsolved, it is a potential limitation for using GANs as a prior for perception reconstruction as the model would never learn to reconstruct the complete space of possible images sufficiently. However as demonstrated in Figure 3 the DCGAN chosen here appears to have learned plenty of the necessary variability.

In conclusion we believe that our method and results present a promising foundation for future extensions. As generative modeling is one topic explored extensively in the machine learning community at the moment, many drawbacks may be solved in the near future. We believe reconstruction of arbitrary visual stimuli during perception, imagination and even dreaming is still a largely under-explored territory of neuroimaging research and will continue to strongly benefit from new advances in the machine learning community.

## 5 Funding

This research was supported by VIDI grant number 639.072.513 of The Netherlands Organization for Scientific Research (NWO).

### 6 Acknowledgements

We would like to express our gratitude to Kendrick Kay, who provided us with an updated `vim-1` dataset, including data for a third subject. We would also like to thank Jonas Kohler and Paul Zhutovsky for discussions.

An example implementation of the method presented here is available on the public GitHub profile of the group: github.com/artcogsys/ganrecon.

## A Supplemental information

### A.1 Training of the deep convolutional GAN

#### A.1.1 DCGAN architecture

The *generator network* consists of one linear and four deconvolutional layers, each followed by batch normalization and rectified linear activation functions (ReLUs). The linear layer takes **z** and maps it to the first deconvolutional layer that expects 512 feature channels. The generator then maps to 256, 128, 64 and 1 feature channels across the deconvolutional layers. Kernel sizes are 4 × 4 and stride is 2 in every deconvolutional layer. The pixel output of the generator is scaled between [−1, 1] by applying tanh to the output values as a final step. Numerical instabilities required additional clipping of the generated pixel values at [−1, 1]. A feature matching loss, using the first discriminator layer, was also added to the generator loss term. This is a common trick: The distance between the desired image feature statistics for real images and the feature statistics for fake images should be minimized. For the `vim-1` DCGAN we manually apply the circular mask used for creating the stimuli at the end of training in order to let the training process focus on the visible area. The *discriminator network* consists of 4 convolutional layers, followed by batch normalization and exponential linear activations (ELUs, (Clevert et al., 2015)). Before the image enters the discriminator handicap Gaussian noise with a standard deviation of 0.15 is added to the input images. This (instance noise) is another common trick, inhibiting discriminator performance to allow the generator to catch up. Except in the initial layer (which had 3×3 kernels) all layers use kernel sizes of 4×4 and a stride of 2. The layers map from 1 to 32, then 64,128 up to 256 feature channels, and are followed by a linear layer mapping all final activations to a single value reflecting the discriminator decision.

#### A.1.2 Training parameters

The latent variable **z** ∈ [−1, 1]^50^ is randomly drawn from a uniform distribution and restricted to a unit hypersphere by normalizing it, in order to embed it in a continuous bounded space without borders. This step facilitates the prediction of **z** in an otherwise unbounded solution to the regression problem. For optimizing the weights of the DCGAN we used the Adam optimizer (Kingma and Ba, 2014) with default parameters (*α* = 0.001, *β*_1_ = 0.9, *β*_2_ = 0.999, *ϵ* = 10^−8^). The learning rate was 10^−4^ for all networks. We applied gradient clipping with a threshold of 10.

#### A.1.3 Training data sets for the three stimulus spaces

The DCGANs for natural images were trained on a downsampled 64 × 64 variant of ImageNet (made available with (Chrabaszcz et al., 2017)) together with the Microsoft COCO dataset^11^. The image size of MS COCO was decreased to 64 x 64 and center-cropped, and images for which this was not possible due to aspect ratio were removed from the training set. Before entering training all images were converted to gray scale and contrast-enhanced, similar to the transformation described in (Kay et al., 2008) (remapping the pixel values to a new range and saturating the bottom and top 1% of the pixel values). The image value range entering training was [−1, 1]. For the `vim-1` GAN, again the circular mask was applied. This resulted in approximately 1.500.000 gray scale natural images used for training in total. Note that DCGAN training would usually also work with a lower amount of training data.

The DCGAN on handwritten characters was trained on (in total) 15.000 examples of B, R, A, I, N and S characters from (Van der Maaten, 2009) and (Schomaker and Vuurpijl, 2000). As the experiment on the BRAINS dataset should focus on a restricted stimulus domain its DCGAN does not require more expressive power.

### A.2 Details on hyperalignment

The common representational space (Haxby et al., 2011) was built using a similar procedure as in (Güçlü and van Gerven, 2015). For each ROI the initial representational space was the data of the subject with the most voxels in this ROI. The representational space was estimated on the training data, and applied on the validation data, projected separately for each subject. The final functional data in the common representational space was then acquired by taking the average over these individual subject projections.

#### A.2.1 Number of voxels per ROI in BRAINS

The two-repetition averaged data from the training and validation set was used for the projection for this data set. It was estimated on the concatenated V1 and V2 data without further separation.

**Table 1:**
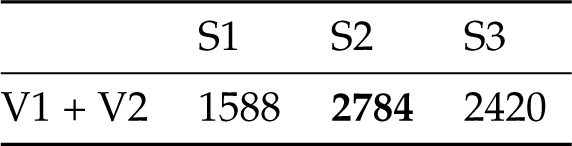
Number of voxels for each subject in the BRAINS data set, which is restricted to V1 and V2. The boldly printed number shows the common representational space dimension.

#### A.2.2 Number of voxels per ROI in vim-1

The original single trial recordings for train and test were used for hyperalignment. These are not publicly available, but could be acquired via personal communication. Due to the large number of voxels, the hyperalignment was done separately for every ROI.

**Table 2:**
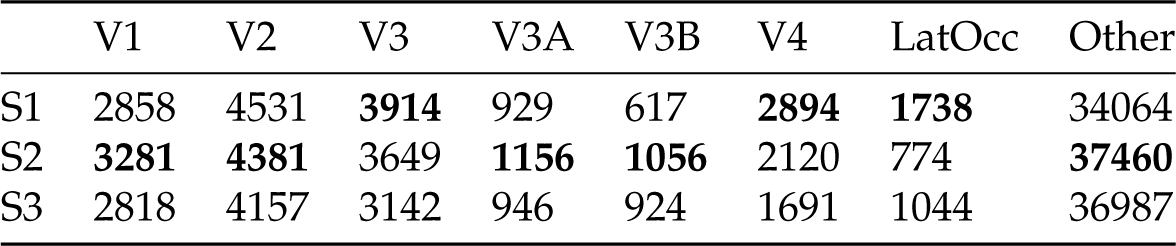
Number of voxels for each subject in the `vim-1` data set. The boldly printed number indicates the respective common representational space dimension for this ROI.

|  | V1 | V2 | V3 | V3A | V3B | V4 | LatOcc | Other |
| --- | --- | --- | --- | --- | --- | --- | --- | --- |
| S1 | 2858 | 4531 | **3914** | 929 | 617 | **2894** | **1738** | 34064 |
| S2 | **3281** | **4381** | 3649 | **1156** | **1056** | 2120 | 774 | **37460** |
| S3 | 2818 | 4157 | 3142 | 946 | 924 | 1691 | 1044 | 36987 |

#### A.2.3 Number of voxels per ROI in Generic Object Decoding

As the voxel dimension was relatively small, as in the `BRAINS` data set we estimated the common representational space at the same time on all voxels. The available ROIs in this data set were V1, V2, V3, V4, LOC, FFA and PPA. The original single trial data was used for all steps.

**Table 3:** Number of voxels for each subject in the `Generic Object Decoding` data set. The boldly printed number indicates the common representational space dimension.

|  | S1 | S2 | S3 | S4 | S5 |
| --- | --- | --- | --- | --- | --- |
| All ROIs | 4466 | 4404 | **4643** | 4133 | 4370 |

## Footnotes

1 https://crcns.org/data-sets/vc/vim-1 (last access May 2017)

2 We focus on Reconstructing gray scale images as our natural images DCGAN learned to generate more structural detail when the color dimension was omitted. However with a more powerfiil GAN variant the method could also be applied for reconstructing color Stimuli.

3 http://brainliner.jp/data/brainliner/Generic_Object_Decoding (last access August 2017)

4 www.pymvpa.com,v2.6.3

5 Our method was initially developed on the Individual Subject basis. This only seemed to lead to more variability in the Reconstruction Quality between Subjects, and we decided to finalize the study on hyperaligned data instead as this made collecting behavioral data and developing the loss function more efficient.

6 A proposed alternative for plain MAE loss would be the MS-SSIM loss (Ridgeway et al., 2015). However, we instead decided to enhance this loss with a series of feature (perceptual) losses.

7 It is not advisable to use MSE between the whole feature matrices as across a convolutional neural network hierarchy for any given image they tend to be sparse. Due to this when using MSE our reconstruction model frequently fell into a local minimum of feature activations with the value 0, which equates blurry images without prominent edges or features.

8 www.chainer.org; Chainer v1.24

9 The influence of omitting the colour information contained in the original stimuli is unclear. Random generations from a DCGAN trained with RGB information missed the structural detail obtained in the grayscale variant (data not shown). When using this RGB DCGAN for reconstructing it was often possible to reconstruct the correct hue for an image and its components. In terms of structure in most cases it was only capable of reconstructing salient blobs however.

10 www.mturk.com

11 www.mscoco.org, described in (Lin et al., 2014) (last access March 2017)

